# Developmental Single-cell transcriptomics in the *Lytechinus variegatus* Sea Urchin Embryo

**DOI:** 10.1101/2020.11.12.380675

**Authors:** Abdull J. Massri, Laura Greenstreet, Anton Afanassiev, Alejandro Berrio Escobar, Gregory M. Wray, Geoffrey Schiebinger, David R. McClay

## Abstract

Here we employed scRNA-seq coupled with computational approaches to examine molecular changes in cells during specification and differentiation. We examined the first 24 hours of development of the sea urchin *Lytechinus variegatus* (*Lv*) with 18 time points during which the embryo develops to the larval stage. Using Waddington-OT, the time points were computationally “stitched” together to calculate developmental trajectories. Skeletogenic cells displayed the expected immediate early divergence while other lineages diverged asynchronously, with many cells retaining an intermediate specification status until late in gastrulation. The *Lv*-scRNA-seq dataset was compared to the developmental Gene Regulatory Network (dGRN) model of specification in *Strongylocentrotus purpuratus* (*Sp*). 79 of 80 genes (98%) in that dGRN are present in the *Lv*-scRNA-seq dataset, and expressed in the correct lineages in which the dGRN circuits operate. Surprisingly, however, many heterochronies in timing of first expression of dGRN genes have evolved between the two species. Replotting the two dGRNs with precise attention to time of expression revealed a number of feedback inputs that likely buffer the dGRNs, allowing them to maintain function in the face of accumulating heterochronies.

**Summary statement:** The early development of the sea urchin embryo was followed using scRNA-seq plus computational methods to trace lineage diversifications. These were matched to gene regulatory network changes over time.

## Introduction

During the earliest stages of development, cells diversify through molecular specification, coordinate their spatial positioning, undergo directed cell rearrangements, and establish the basic body plan. Later development includes a massive increase in body mass while cells specialize into tissues and organs. This sequence occurs rapidly, and with high fidelity. Errors are rare, and often catastrophic when they occur. At the same time, development is considered to be one of the major arenas for evolutionary change, thus raising the question of how development operates within a constrained, high-fidelity system and yet is subject to evolutionary change.

Emerging technologies provide increased leverage to address this question and to explore the complex nature of development. In particular, recent progress has been strongly augmented by high-throughput single-cell measurement technologies like single cell RNA sequencing (scRNA-seq), combined with analytical approaches to infer developmental trajectories in mouse and zebrafish embryos (Farrell et al., 2018; Schiebinger et al., 2019), as well in a growing number of embryos, tissues, and disease states (Cao et al., 2017; Fincher et al., 2018; Han et al., 2018; Karaiskos et al., 2017; Plass et al., 2018; Wagner et al., 2018). Due to the changing nature of development, and the cell’s destruction during scRNA-seq protocols, it is only possible to gain a snapshot of a single cell’s transcriptome. To measure temporal changes with scRNA-seq, a compromise solution to the snapshot issue is to capture cells at a series of timepoints that are relatively close together, then computationally stitch them together to infer a continuous sequence. It is important to assess whether the spacing of the timepoints is sufficiently close to allow the temporal change to be of value for mechanistic inference. If so, an ability to follow the lineage sequence enables the reconstruction of an atlas of transient cell states over time.

Among the questions in development that are a work in progress is that of cell fate specification. For two decades developmental Gene Regulatory Networks (dGRNs) that direct cells toward their fates have been studied in many embryonic systems (Peter and Davidson, 2015). dGRNs of various model embryos are inferred or empirically established through iterative perturbations of candidate signals and transcription factors. These GRN models and the accompanying advent of systems biology revealed a number of molecular devices that govern network circuitry during the progression of cells through specification and commitment. Feed-forward and feedback circuits are among the transcriptional devices that advance and stabilize GRN states (Yeger-Lotem et al., 2004). These and many other regulatory mechanisms continue to be uncovered in many embryonic systems (Zhou et al., 2018). The architectural features of network circuits provide the functionality that lead to a cell fate and ultimately to tissue assemblies. Changes in these circuits are hypothesized to be drivers of evolutionary diversification (Erkenbrack and Davidson, 2015; Erwin and Davidson, 2009; Hinman and Davidson, 2007; Hinman et al., 2003; Israel et al., 2016). Thus, exploration of GRNs by a powerful technology such as scRNA-seq, promises new insights to our understanding of how embryonic systems work, and evolve.

To begin using scRNA-seq for inference into regulatory analysis, a number of steps must be followed. A necessary step is to determine whether the depth of sequencing using this technology is sufficient to detect expression of the transcription factors and signals that constitute the specification networks. To explore that question we chose to analyze the sea urchin dGRN. Over the past two decades, a number of investigators experimentally assembled a dGRN of sea urchin embryo specification. Many perturbations of transcription factors and signals from multiple laboratories revealed a detailed network model that operates from fertilization until gastrulation (Davidson et al., 2002; McClay et al., 2020; Peter and Davidson, 2015). Recently, scRNA-seq was used to compile the first atlas of development for *Strongylocentrotus purpuratus* (*Sp*) (Foster et al., 2020), and that approach was further used to explore the development of pigment cells in *Sp* (Perillo et al., 2020). Given the extensive knowledge of dGRNs in the sea urchin embryo, and taking advantage of the methods for scRNA-seq developed for this model embryo {Massri, 2021; Perillo et al., 2020), our first goal was to establish whether the molecules in dGRNs were “visible” to scRNA-seq inquiries. We find that more than 98% of the molecules in the sea urchin dGRN are present in our scRNA-seq database, both in the lineages expected and at the times the dGRN models indicate those signals and transcription factors are present to guide specification. This gave us license to explore the dGRN further. We explored the cell lineages and their accompanying dGRNs using a recently developed computational tool “Waddington-OT” (Schiebinger et al., 2019). We also developed novel computational tools utilizing barycentric coordinates to follow lineage progressions and dGRN changes over time. The details of these dGRN and lineage findings are provided following a series of quality checks. These established that the atlas of cells produced is of high quality and accurately reflects cell lineage progression from zygote to larva.

The ability to follow each reconstructed lineage using scRNA-seq, supplied by thousands of gene expression patterns, provided an opportunity to explore novel properties of cell diversification. In some cases, the divergence was simple as suggested by a seminal asymmetry event. However, other lineage separations were gradual, implying a much more complex series of events necessary to finally resolve a cell type. Further, a comparison between *Lv* and *Sp* revealed a number of temporal changes in dGRN dynamics that have evolved during the 40 – 50 million years since the common ancestor of these species. Despite those many changes, analyses revealed new stabilizing features of the dGRNs that likely contributed to the high degree of conservation of the dGRNs between the two species.

## Results

### Atlas of sea urchin development from fertilization to larval stage

The sea urchin embryo is a basal deuterostome that has long been a model for studies of early development. Its development to the larval stage is rapid, occurring in *Lv* within 24 hours. To map known GRN genes, and to identify novel candidates during this critical time period of embryonic development, we performed a scRNA-seq analysis using the 10x Genomics system. This standardized and reproducible advanced scRNA-seq platform yields relatively deep coverage of RNA expression in single cells at a cost that is not prohibitive (Massri et al., 2021; Perillo et al., 2020). Eighteen timepoints were collected, initially at hourly intervals and then during later stages at two to four hour intervals. Rather than process the same number of cells for each timepoint, we increased the cell number with increasing stages to cover the increased developmental complexity. To generate the number of embryos and biological diversity needed for the study, eggs from six females were fertilized by sperm from one male *Lv* sea urchin. At each time point approximately 10^ embryos were dissociated to single cells and then fixed, using a published protocol we adapted (Alles et al., 2017; Chen et al., 2018; Juliano et al., 2014; Massri et al., 2021; McClay, 1986). Each time point separated cells from the embryo under conditions that minimally affected the mRNA expressed in each cell. The dissociated cells were immediately fixed under conditions that retained more than 95% of the resident mRNA per cell. Fixed cells were then rehydrated and counted before cell encapsulation and library preparation. Following library preparation, samples were sequenced at >50,000 reads/cell to ensure read depth and coverage (Haque et al., 2017; Svensson et al., 2017; Ziegenhain et al., 2017).

According to a standard analysis pipeline (Massri et al., 2021), we mapped reads to the Lv3.0 genome (Davidson et al., 2020), with the 10x Cellranger pipeline and filtered out low quality cells using Seurat (Butler et al., 2018; Stuart et al., 2019). In total 50,935 cells remained and were used for the analysis. We obtained a matrix of raw gene expression counts for each sample, which was normalized using SCTransform, a regularized negative binomial regression method that stabilizes variance across samples (Hafemeister and Satija, 2019), then visualized using UMAP (Uniform Manifold Approximation and Projection) (Becht et al., 2018). **Fig. 1** shows the requisite overview UMAP visualizations of these 50,935 cells. Diagrams of embryos at several stages of development (**Fig. 1A),** relate developmental states to hours samples were collected. The UMAP plot in **Fig. 1B** is colored according to 16 embryonic cell types that emerge during that 24 hr period. The colors match the cell types shown in the embryo diagrams of **Fig. 1A. Fig. 1C** shows the same UMAP colored to show the hours of collection. These colors match the dot colors shown in the time course diagram. Graph-based louvain clustering, implemented in Seurat, was used to obtain 63 distinct clusters of cells, and these were annotated using dGRN marker genes (Davidson et al., 2002; McClay et al., 2020; Peter and Davidson, 2015), to provide a preliminary identification of the embryonic lineages. **Fig S1A** shows the major annotated lineages plotted over the 18 time points and **Fig. S1B** shows the 63 clusters. The dynamics of the temporal progression of stages is shown in **Movie S1.** The cells in the movie are colored to show the initiation, divergence, and progression of lineages as a function of time.

**Fig. 1:**
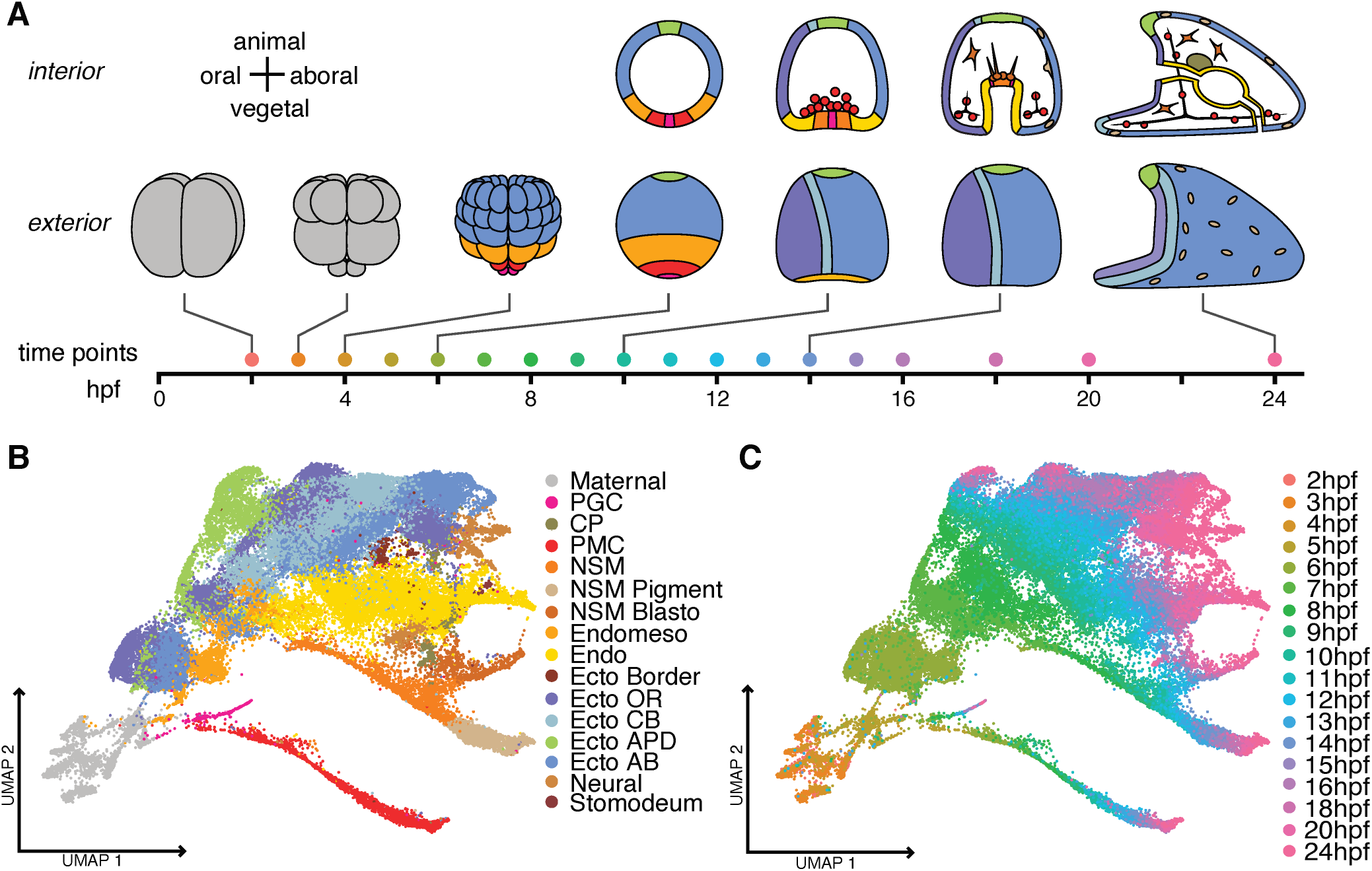
Temporal atlas of scRNA-seq profiles during development of the sea urchin, *Lytechinus variegatus.* **A.** A time-line of development over 24 hours with representative stages of development illustrated. Colors represent lineages as in Fig.1B: ectoderm (blue), oral ectoderm (dark blue), aboral ectoderm (light blue), skeletogenic mesoderm (red), primordial germ cells (pink), non-skeletogenic mesoderm (orange), pigment cells (beige), endoderm, (yellow), ciliary band (violet), animal pole domain (green), early cleavage (gray). The dots on time points are colored according to the time points as in Fig. 1C. **B.** UMAP plot mapped according to lineage domains. Sixty three clusters were identified and grouped into 16 domains representing the lineages present at 24 hpf. The colors are the same as for the embryos diagrammed in Fig. 1A. **C.** The UMAP plot colored according to time of development showing the position of the 18 timepoints collected during the first 24 hrs post fertilization (hpf). The colors along the time-line are the same as those in the UMAP plot. PGC = primordial germ cells; CP = coelomic pouch; PMC = primary mesenchyme cell (also called skeletogenic cells); NSM = non-skeletogenic mesenchyme; Blasto – blastocoelar cells; Endo = endoderm; Ecto = ectoderm; Border = ectoderm at the border between ectoderm and endoderm; OR = oral; CB = ciliary band; APD = animal pole domain; AB = aboral.

The structure of the UMAP recapitulates the basic events of sea urchin development. It has long been established that the skeletogenic (or primary mesenchyme cells (PMCs)), and primordial germ cells (PGCs) branch from specification of the other lineages as early as the 4^th^ and 5^th^ cleavages by two successive unequal cleavages (McClay, 2011). The UMAP displays that separation, and provides a continuous track of the two lineages as they are specified over time. In addition, the endomesoderm later splits into the two canonical germ layers: endoderm and mesoderm. This is once again reflected in the UMAP plot since at 5 hours post fertilization (hpf), cells of both germ layers are clustered together indicating that the cells still have not yet diversified. By 7-8 hpf the endoderm and mesoderm cells have separated, and later each further subdivides as diversification within the germ layers occurs. Meanwhile, specification of the ectoderm occurs leading to regional anterior-posterior, dorsal-ventral, and right-left differences in this lineage (**Fig. 1B**; **Fig. S1**).

As part of the quality control of the analysis, we asked whether the pipeline from embryo to sequencing output introduced any bias in the relative abundance of cells from each lineage. Based on earlier lineage analyses of embryos (Davidson et al., 1998; Logan and McClay, 1999; Martik and McClay, 2017), the distribution of cells allocated to ectoderm, mesoderm and endoderm appeared not to be biased by over- or under-selection once the lineages could be identified by genes expressed at 5 - 6 hpf (**Fig. S2**). Since allocation of cells to lineages was based on expression of all genes in a cell, prior to five hours the prevalence of maternal transcripts distributed to all cells dominated and prevented the lineages from being separately distinguished. At the 5 hour time point, lineage markers had emerged providing an imperfect approximation of lineages, and from 6 hpf until the larval stage, the expected number of cells for each lineage was present. At 6 hpf there were no cells in the non-skeletal mesoderm because at that time point this lineage had yet to emerge from the endomesoderm (shown in the endoderm lineage only).

We found that even relatively rare cell populations could be detected. Normally, four Primordial Germ Cells (PGCs) arise at 5^th^ cleavage, later divide once, and at gastrulation migrate to the coelomic pouches without dividing further (Campanale et al., 2014; Voronina et al., 2008; Martik and McClay, 2015). Despite contributing a maximum of 8 cells per embryo, we detected PGCs at each time point, providing a continuous record of that lineage (**Fig. 1**, **Fig. S1**). Another rare cell type in the embryo are the cells that become serotonergic neurons. In *Lv* at the pluteus larval stage only 4-5 such cells are detected (Slota and McClay, 2018). Again, that rare cell type was detected (**Fig. S1B**), giving us confidence that our developmental reconstruction reflected a detailed and accurate representation of each lineage’s transcriptional history over the first 24 hours of development.

### Detection of molecules contributing to developmental gene regulatory networks

We next sought to determine whether the samples were sequenced deeply enough to detect the transcription factors known to participate in specification of the lineages of sea urchin development. The data revealed that the scRNA-seq analysis faithfully reflected expression of known transcription factors in the network of each previously characterized lineage. We analyzed the expression of 80 transcription factors and signals from the endomesodermal and ectodermal dGRN models (Li et al., 2013; Peter and Davidson, 2015). We first determined that all but one gene in the dGRN models are present in the relevant lineages indicating that the scRNA-seq database can provide information on 98% of the known dGRN genes (the missing gene is *twist,* a short gene expressed at very low levels). 52 of those 80 genes are plotted in a quantitative dotplot shown in **Fig. 2**, and all 80 are shown in UMAP distributions in **Fig. S3A** and **S3B**. For some transcription factors known to be expressed at low levels, there was an expected reduction in the proportion of the cells of a lineage in which that transcription factor was detected by scRNA-seq. For example, *Snail* is a gene known to be lowly expressed where ever it is found (Cano et al., 2000; Wu and McClay, 2007; Materna et al., 2010). Predictably, the percentage of cells in which a *snail* sequence was reported (**Fig. S3B**), was small relative to detection of other transcription factors that were relatively abundant as quantified in **Fig. 2**. Nevertheless, even a gene such as *snail* could be followed in the lineage trajectories (**Fig. S3B**). We conclude that the scRNA-seq database has the sensitivity to examine the spatial and temporal patterns of expression of the vast majority of transcription factors and genes used in specification of embryonic cells, including, we suspect, transcription factors that contribute but are not yet included in the dGRNs.

**Fig. 2:**
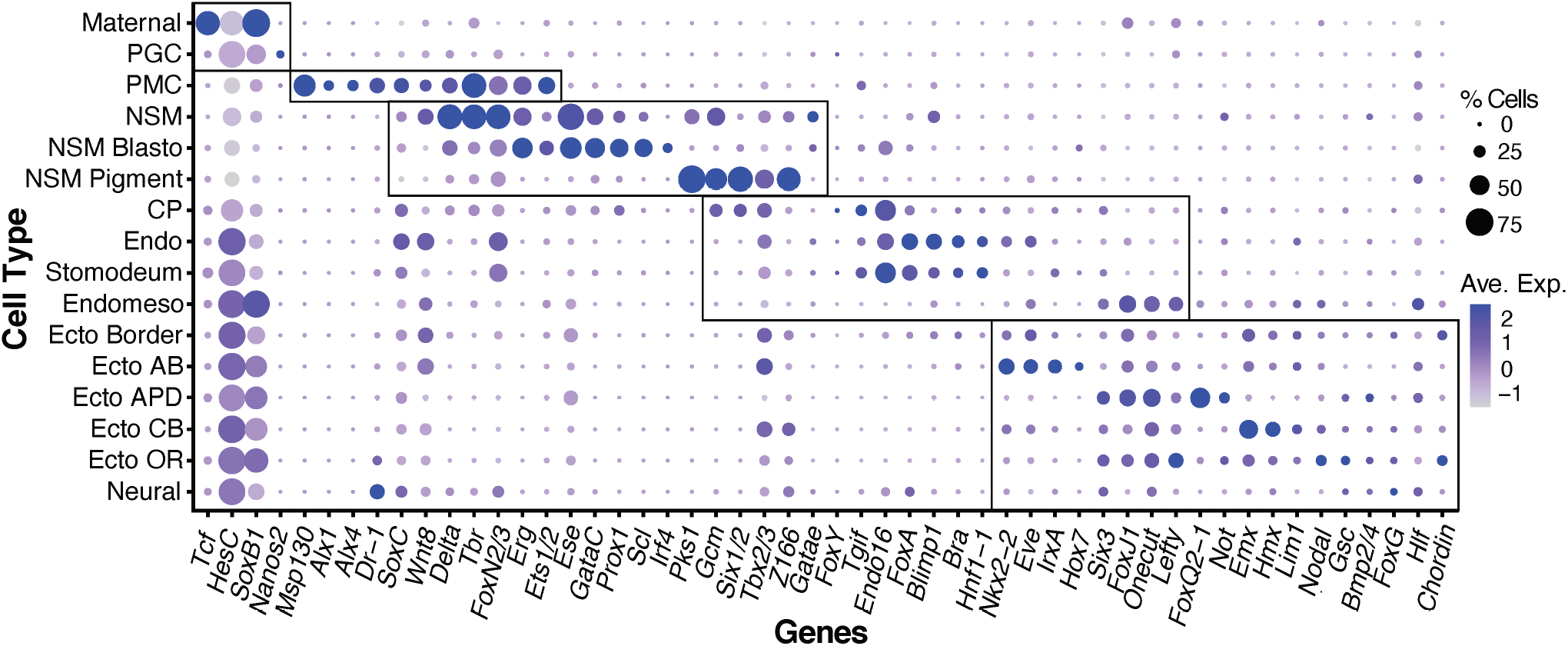
Dotplot of genes expressed in the sea urchin developmental Gene Regulatory Network (dGRN). 52 of the more than 80 genes expressed in the dGRN models (Ben-Tabou de-Leon et al., 2013; Li et al., 2013; Peter and Davidson, 2015) were plotted according to expression in the clusters listed on the Y axis. The dots of each shows relative level of expression in that cluster as well as the percentage of cells in that cluster that express that gene indicated by size of the dot. Boxes surround the clusters assigned to PGC, PMC, NSM, Endoderm and Ectoderm. Some genes are expressed outside the boxes. Those genes are expressed in more than one lineage, sometimes at the same time and in other cases at different times.

### Inferring developmental trajectories with Waddington-OT

We next sought to infer developmental trajectories and analyze the diversification of the various lineages, with the ultimate goal of identifying transcription factors regulating these diversification events. We applied Waddington-OT methods to connect the scRNA-seq data from different time-points and infer developmental trajectories (Schiebinger et al., 2019). Optimal transport connects cells sampled at one time point to their putative descendants at the next time-point in a way that minimizes differences in expression over all genes. The algorithm requires as input an estimate of each cell’s proliferative ability, which it uses to allocate each cell a certain number of descendants. Based on the concordance between observed and expected changes in abundances of each of the five primary lineages (**Fig. S2**), cell proliferation rates were assigned, and we ran Waddington-OT with the default parameter values (see Methods for details). The resulting output is a time-course of transport matrices connecting each pair of time-points. These matrices have a row for each early cell and a column for each late cell, and the i,j entry is interpreted as the number of descendant’s cell i would have of type j at the later time-point.

We tested the quality of our inferred trajectories by demonstrating that we could interpolate the distribution of cells at held-out time points, as described in (Schiebinger et al., 2019). This was important as it established that our temporal resolution was sufficient to accurately stitch together cells from adjacent timepoints (**Fig. S4A**). In other tests this result was robust to downsampling cells or reads, moderate changes in parameter values away from the default settings, or to moderate perturbations to growth rates (**Fig. S4B-F**).

### Visualizing lineage diversifications

To visualize the divergence of various lineages, the transport matrices compute, for any cell from an early time-point, the proportion of descendants that obtain various fates at later time-points. We developed a simple way to visualize these ‘fate probabilities’ for triples (or quadruples) of lineages in a triangular (or tetrahedral) plot (**Fig. 3**; **Movie S1**). The visualization employs barycentric coordinates to represent k-dimensional probability vectors in k-1-dimensional space. For example, suppose we wish to visualize the cells from 8 hpf according to their probability of giving rise to an ectodermal, endodermal, or ‘Other’ descendant at 20 hpf. We identify a vertex of the triangle for each of these possible fates, and position the cells according to their relative probabilities. Cells perfectly fated to obtain a single fate are positioned exactly at the corresponding vertex, while cells with indeterminate fates are positioned in the interior of the triangle. The very center of the triangle corresponds to cells that are equally likely to choose any of the three fates, and cells along an edge have zero chance of reaching the opposite vertex. **Fig. 3** illustrates examples of how selected pairs of lineages diverge in *Lv* over time. **Movie S1** shows a tetrahedron visualization of the divergence of four fates simultaneously, in parallel with the progression of those same cells along the UMAP over time. These visualizations allow us to easily identify groups of cells with specific fate probabilities as they migrate towards the corners over time. Moreover, expression of individual genes or gene signatures can be visualized in these barycentric coordinates much in the same way as they are on a UMAP plot.

**Fig. 3:**
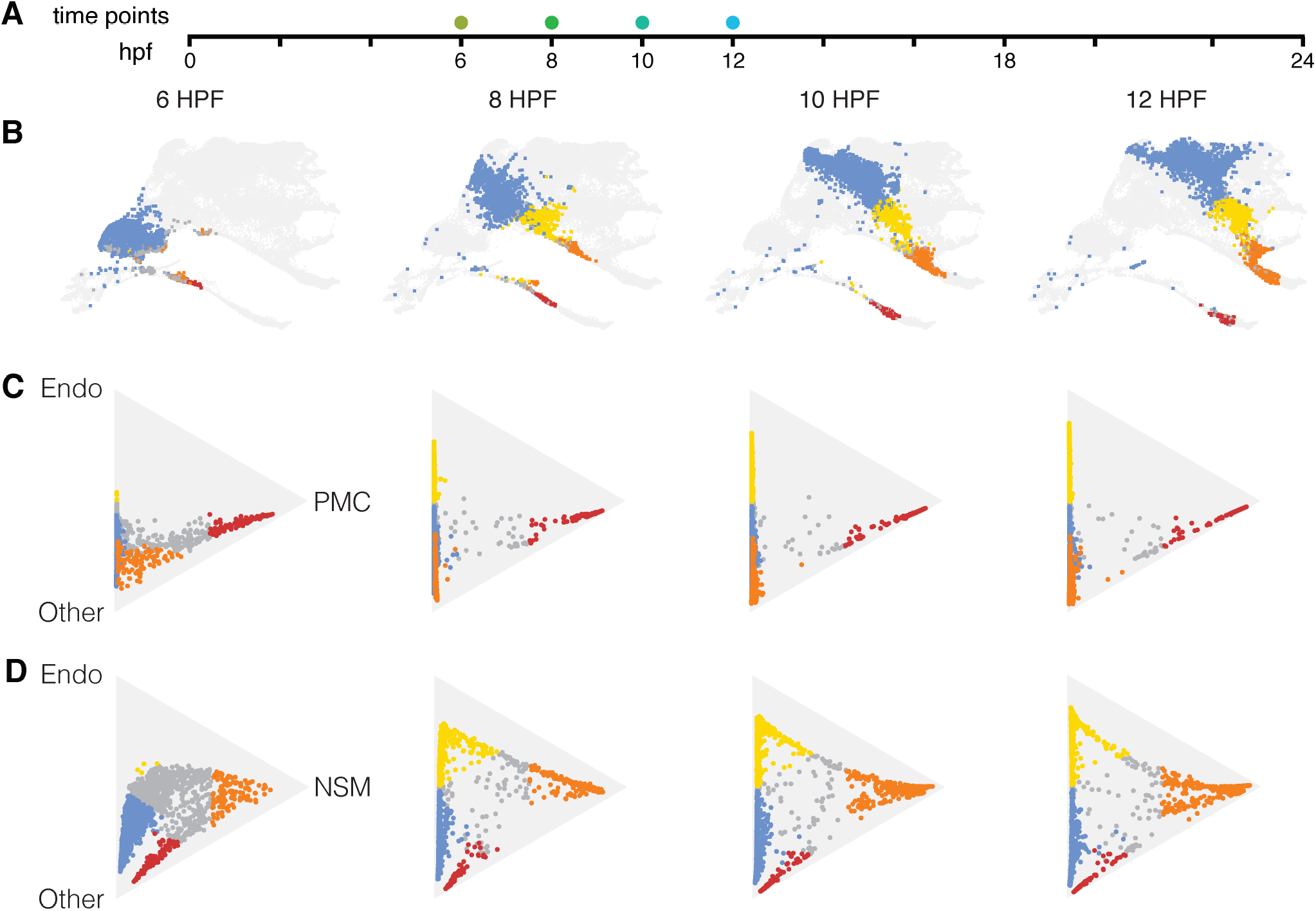
Diversification of Cell Lineages computed using the Waddington-OT optimal transport method. (Schiebinger et al., 2019). **A.** Time-line represented by the plots with dots to indicate the times shown. The 6 hpf time point is the blastula stage, 8 hpf = hatched blastula; 10 hpf = mesenchyme blastula; and 12 hpf = early gastrula stages. **B.** UMAPs at those four time points showing presumptive ectoderm (blue), endoderm (yellow), NSM (orange) and PMCs (red). **C.** Triangle plots show progression of lineages at different time points, with vertices designated Endoderm, PMC, and “other” to represent all other lineages. D. Triangle plots with vertices designated Endoderm, NSM, and Other to represent all other lineages. The colored cells in the triangle plots are those cells with at least a 60% probability of becoming one of the colored lineages represented. Cells that have not reached a 60% probability toward any of those fates are colored gray. Cells that hug a side of the triangle have low to no chance of becoming a cell on the opposite vertex. Cells in the exact middle of the triangle have an equal chance of becoming a cell of any type. Two major patterns are shown. On the top row PMCs are committed to one side of the triangle from the earliest time point onward. On the bottom row the NSM and Endoderm lineages are committed later with many gray cells entirely traversing the triangle to the opposite side with an extended delay in their commitment toward endoderm or NSM.

### Embryonic lineage diversification differences as seen using the transport matrices

Often textbooks in development indicate that cell diversification begins when a molecular asymmetry occurs before or just after cell division so that two progeny diverge toward alternative fates. The simplest of models describes a factor’s asymmetric distribution driving the separation. Once an asymmetry is in place, the two lineages are modeled as going their separate ways from that time forward. An example of this type of divergence in the sea urchin occurs at the fifth cleavage at the vegetal pole when skeletogenic cells diverge from the fates of other cells. During that 30 min cleavage cycle, skeletogenic cells activate *pmar1,* a repressor that represses a repressor, *hesC* (Revilla-i-Domingo et al., 2007). All other lineages in the embryo fail to activate *pmar1.* As a consequence, *hesC* is activated in all non-skeletogenic cells during that 30 minute interval. This results in repression of the skeletogenic fate in those lineages. **Fig. 3C** shows the consequence of that diversification. At all time points, cells destined to become PMCs follow one side of the triangle, the side leading to the PMC fate.

However, cell fate decisions are not always so simple and clear. **Fig. 3D** shows the diversification of the endomesoderm. There, the endomesoderm ancestors diverge as a wave of cells spanning the distance across the triangle and that wave continues to spread through time until cells at later time points reach their eventual fates. In contrast to the skeletogenic cells in **Fig. 3C**, many cells tend to remain in an intermediate state until late in specification. Then, much later, as the cells near the time of differentiation, the wave parts, sending cells toward one of the two fates (see Movie S1 for an animated depiction of this sequence). This outcome indicates that the fate decision is both non-uniform temporally, and often quite late, even though the cells are continually specified (the wave progression). Further, the broad distribution of cells in the wave between the two final fate points suggests that the specification is variable in a lineage for much of the time allowing for many of the cells to go either way until late in the temporal progression. We will return to this intermediate state below, but first it is necessary to assess the concordance between the scRNA-seq temporal profiles and the known dGRNs.

### Matching cell diversification to developmental GRNs

Since we had established that 98% of the transcription factors and signals in the dGRN are present in the scRNA-seq database, we wanted to determine how well the expression of those genes matched the lineages established with the optimal transport method. If the plots of cell populations over time reveal when fates of the cells are decided, as inferred above, the Waddington-OT platform should identify populations of cells that correlate with the dGRN states established experimentally. If that were the case, cells in specification space should reflect the dGRN progression. To test that prediction we examined collections of cells that had diverted toward their final fate by a defined probability. **Fig. 4** shows two populations, one predicted to become skeletogenic cells, the other, endoderm. The red cells in **Fig. 4A** are predicted to become skeletogenic cells with better than 60% probability (at 6 hpf), while **Fig. 4B** indicates at 9 hpf, yellow cells predicted to become endoderm with a 60% probability. The gray cells in each triangle of **Fig. 4A and B** are all “other” cells that are not projected (at the 60% level) to become PMCs or endoderm respectively, though some of those cells may have reached the 60% threshold for an alternative fate. Next to the triangle plots are the position of those same cells in the UMAP plots.

**Fig. 4:**
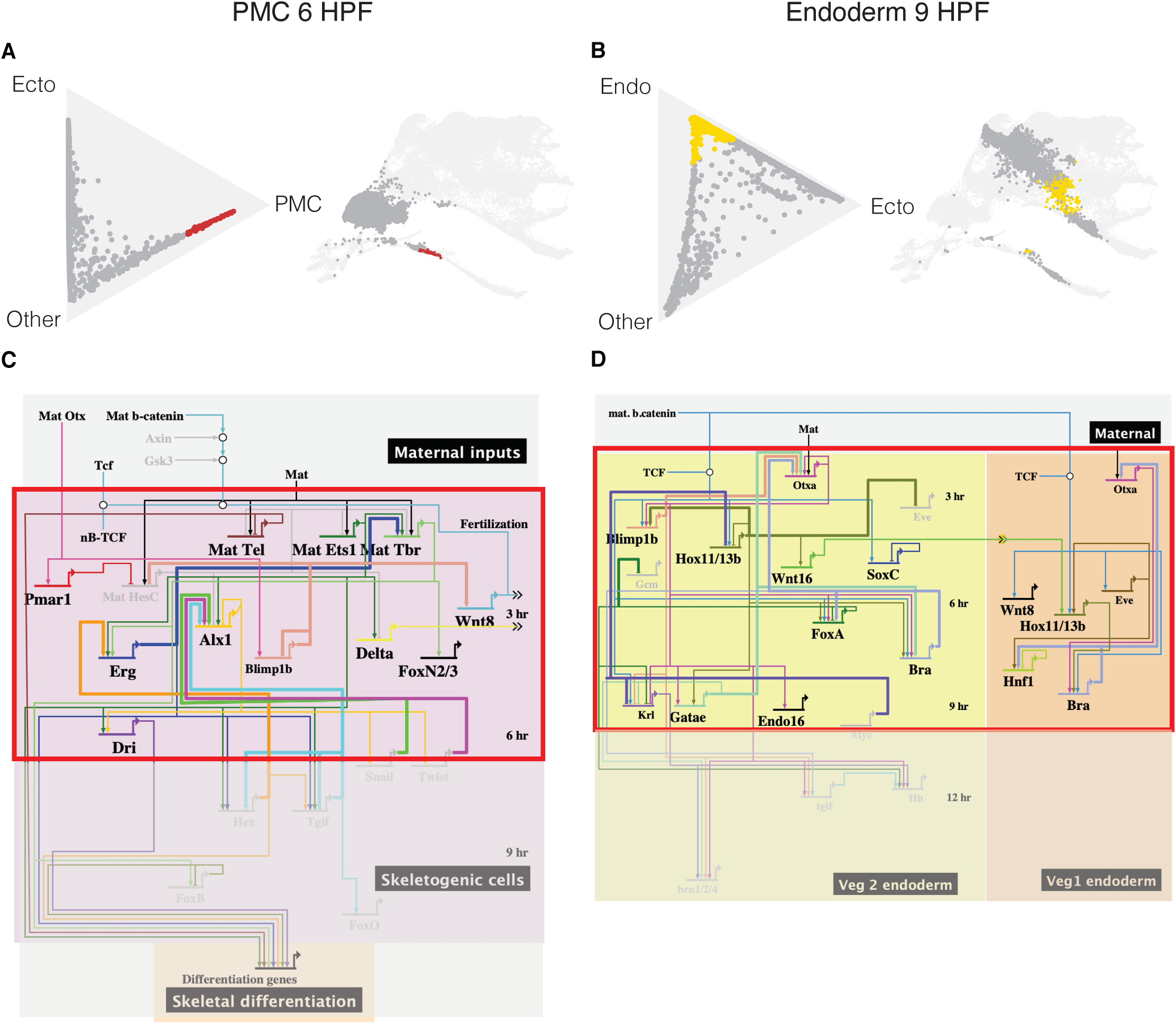
Predictive ability of Waddington-OT plots relative to published dGRNs. **A.** Future PMCs at 6 hpf. The cells in red are those that have >60% probability of becoming PMCs while those in gray either have not reached that level of probability or are in the process of being specified toward other fates. The position of those cells is also shown on the UMAP plot. **B.** Future endoderm seen at 9hpf. In yellow (>60% probability). All other cells are shown in gray. **C.** A dGRN of PMCs with the boxed in area showing genes expressed at 6 hpf. Genes in bold letters are those genes in the top 200 list of genes when the red vs gray genes are compared from the 6 hpf time point in A. **D.** dGRN of endoderm at 9hpf. Shown in the box are all endoderm dGRN genes expressed at 9 hpf. Genes in bold are those in the top 200 list of genes at that time in the endoderm when the presumptive endoderm cells were compared with all other cells in the embryo at that time.

**Figs. 4C and 4D** show BioTapestry (Longabaugh et al., 2009) dGRN models of skeletogenic cells and endoderm cells at the same times as shown on the Triangle and UMAP plots. To ask how well the red skeletogenic or yellow endodermal cell populations matched the dGRNs at those times, the ratios of gene expression between red:grey and yellow:grey were determined and a list of the 200 genes with the highest relative ratios was established. The idea behind the ratios was that if the cells predicted to become skeletogenic cells accurately reflected specification, those ratios should include dGRN signature genes for that fate with a high ratio relative to alternative fates. We went through the top 200 lists at several time points to ask how many of the genes in the skeletogenic or endoderm dGRNs, at the times in question, were present in the top 200 ratio lists (**Tables S1-S4**). **Figs. 4C and 4D** show the match between dGRN state at the time points in question, and the genes on the lists. The BioTapestry models show in bold, genes that are in the top 200 lists. Approximately 71% and 73% of the transcription factors, (modeled at the 6 hpf and 9 hpf times) are present in the top 200 gene lists for the two cell types. This high level of concordance suggests that the optimal transport approach is useful for dGRN inference, and therefore, that other transcription factors that have ratios highly favoring a selected population at this time point are excellent candidates for inclusion in the dGRN model, and should be tested. The remaining genes in the two dGRNs were in the total lists of genes in cells of the two cell types but were not in the top 200 list. This is because the top 200, based on ratio of expression, did not include broadly expressed genes (ß-catenin, TCF, Otxa), or dGRN genes that were also expressed at a high level in another tissue at that time (Blimp1, eve). Thus, while the ratio comparisons do not perfectly predict GRN assignments, they provide a very good predictor of tissue specific genes, and since those lists include large numbers of genes with unknown function in the embryo, they also provide an excellent list of candidate genes for future study.

### Uncommitted “intermediate” cells in the triangle plots express multiple gene signatures

To test the hypothesis that the intermediate cells of **Fig. 3** could be directed toward either of the two nearest fates, we developed a “gene signature” of each cell type using the well described developmental gene regulatory network (dGRN) for the sea urchin as our guide. The prediction was that if the cells were, in fact, intermediate, they should express members of both endoderm and NSM dGRNs. Using signatures of mesoderm and endoderm dGRN transcription factors that drive specification of mesoderm or endoderm (**Fig. S5**), we computationally sampled the specification state of all cells in the middle of the triangles. Cells were colored according to their expressed signatures as they progressed through specification. **Fig. 5** shows the results of that experiment. If a cell expressed only one of the two signatures it tended toward a dark blue color and hugged the edge of the triangle leading to one of the two fates (or if neither signature, it was closer to the “other” fates, i.e. ectoderm and PMCs). The cells in the middle and the cells in the “wave”, by contrast, were colored green or yellow-green or yellow, indicating that both signatures were present (pure yellow indicated a 1:1 ratio of signature genes expressed in a cell). These data support the hypothesis that in separation of endomesoderm into mesoderm and endoderm, some cells proceed early toward a distinct fate, but other cells express transcription factors of both fates for an extended time, and only commit toward one or the other fate with an extended delay. It should be noted that the position of the cells is not based just on expression of the signature genes. That position, as with all the triangle plots, is based on expression of all genes of a cell and the position is established according to the optimal transport method described above. The coloring of the cells in **Fig. 5**, however, is based on the expression of signature genes in cells at those positions. To further test the hypothesis, we randomly sampled individual cells at different positions along the “wave” of intermediate cells to determine which genes of the two signatures were present. Did they simply retain a specification state more like the earlier endomesodermal ancestral state, or, did they express some transcription factors that normally appear once the mesoderm and endoderm diverge? **Fig. S5** shows the results of seven sampled cells (labeled pink). In 6 of the 7 cells signature transcription factors or markers of both mesoderm and endoderm were present. Cells halfway between the two fates tended to express both signatures equally. Cells closer to one of the two vertices tended to express a larger number of the signature genes closer to the nearby vertex than the other. One cell, the cell closest to the mesoderm vertex, expressed only mesoderm signature genes.

**Fig. 5:**
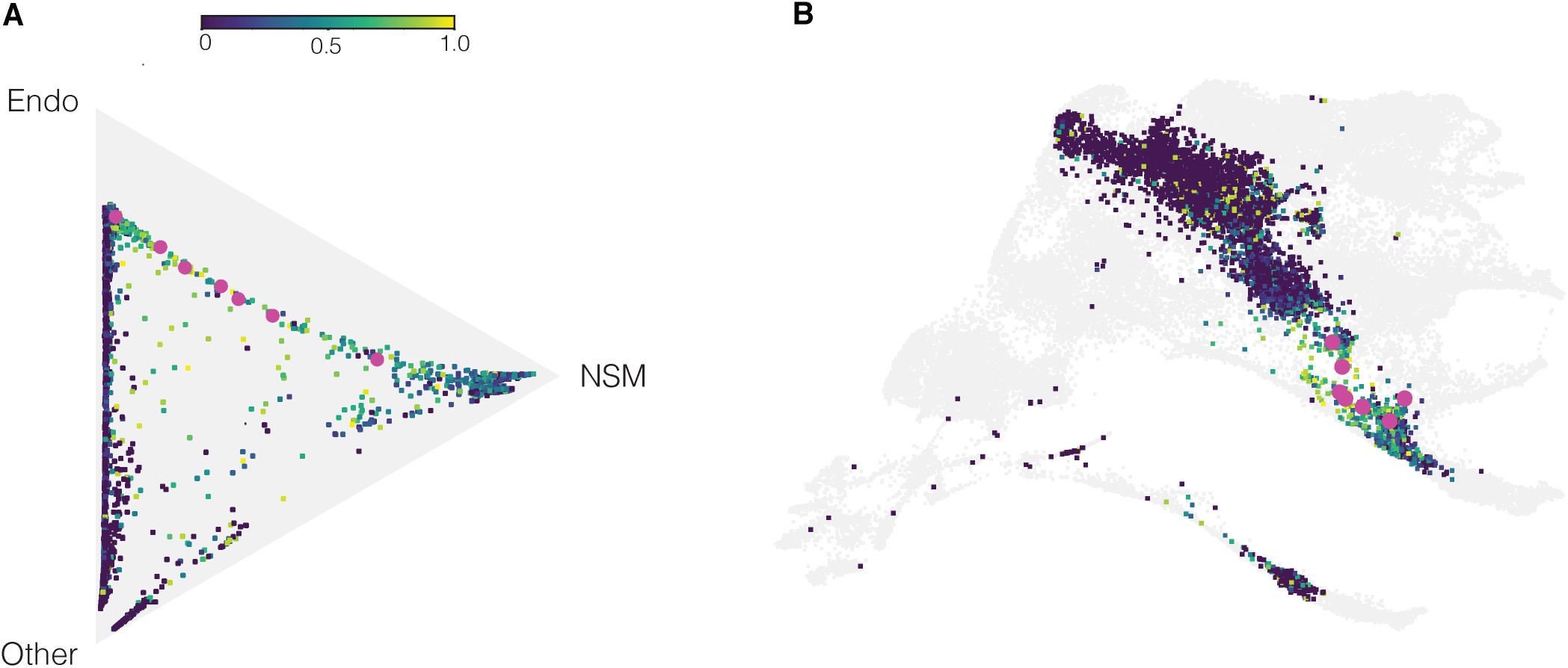
“Intermediate” cell states defined by dGRN gene signatures. **A.** In this plot a group of signature genes expressed by either endoderm or NSM were chosen using the same group of cells shown in Fig. 4B. In this triangle cells that are dark blue are committed to either endoderm or NSM based on expression either of endoderm signature genes only, mesoderm signature genes only, or “other” genes. Cells that are intermediate and express both endoderm and NSM signature genes are colored from yellow (1:1 endoderm;NSM expression) to ratios that trend toward one or the other profile. To ask which signature genes are expressed in samples along the wave of intermediate cells we sampled 7 cells (in pink). **Fig. S5** shows the signature genes expressed in each of these cells. Note: the position of each cell is based on Waddington-OT which uses an algorithm based on all genes expressed by each cell so the signature genes from the dGRNs are but a few of the more than 1000 total genes expressed by each cell, but those transcription factors in the dGRNs likely have a major impact by controlling expression of many of the total number of genes per cell given their regulatory role in development. **B.** The data from the triangle plot shown on the UMAP plot, including the pink cells.

This observation is important as it illustrates that many cells of the mesoderm and endoderm spend extended time in an intermediate specification state before they become committed to one or the other fate. That decision occurs not as a consequence of a single event, as was seen with skeletogenic cells, but likely as a consequence of multiple inputs. Further, we sampled plots as late as 16 hpf and there were still intermediate cells expressing genes of both signatures, though by that time most cells had moved to a likely committed state.

### Timing of gene expression compared between two species

We next turned to a temporal analysis of transcription factor expression in the dGRN. Previously, a number of molecular heterochronies (changes in the relative timing of gene expression) were observed between distantly related sea urchins (Wray and McClay, 1988). A more recent analysis revealed that the skeletogenic lineage of those two species had much in common, though there was some rewiring of the network nodes (Erkenbrack and Davidson, 2015). With a vastly improved temporal resolution now available, we examined the timing of gene expression between *Lytechinus variegatus* (*Lv*) and a relatively closely related species, *Strongylocentrotus purpuratus* (*Sp*). A published high-resolution time-course of gene expression is available for most of the genes in the *Sp* dGRN generated using the nanostring platform (Materna et al., 2010). We adapted the Waddington-OT approach to determine timing of first expression of dGRN genes along lineage trajectories (Schiebinger et al., 2019) (see methods for details).

Transcription factors begin to function well before the steady state of their expression is reached (Bolouri and Davidson, 2003). Thus, a measure of first expression of a transcription factor provides an approximation of when it is deployed in a dGRN (assuming a built-in lag is present). Accordingly, we recorded the earliest expression for 49 of the transcription factors and signals used in early specification and included in the dGRNs for *Lv* and *Sp*. The scRNA-seq approach provided a precision that was unavailable to the earlier nanostring analysis (Materna et al., 2010), because the scRNA-seq profiles allowed us to track individual lineages (**Figs. S1,S3**), while the *Sp* nanostring approach analyzed expression in whole embryos. Thus, if more than one lineage expressed a gene, the nanostring analysis was unable to distinguish from the data which cells were being measured. Fortunately for that analysis, many of the dGRN genes are expressed in only one lineage (as revealed by in situ RNA hybridization) during the time frame covered by the dGRNs. With the caveat that not all the data for *Sp* genes is reported in a lineage specific manner, we compared the timing of expression between the two species where a clear comparison was possible. **Fig. 6** and **Table S5** show the time of first expression for 49 genes in the dGRNs that could be compared between the two species. The rate of development for *Lv* is approximately twice as fast as *Sp* since *Lv* is cultured at 23°C while *Sp* is cultured at 15°C. The two-fold timing difference contributes to rate of cleavage, arrival at canonical stages, and we expected that the same would be true for the timing of first expression of GRN genes. Thus, if a gene in *Lv* was expressed at 3 hpf, the expectation was that the same gene should be expressed at 6 hpf in *Sp.* A variance of two or more hours from that expectation was scored as a heterochrony.

**Fig. 6:**
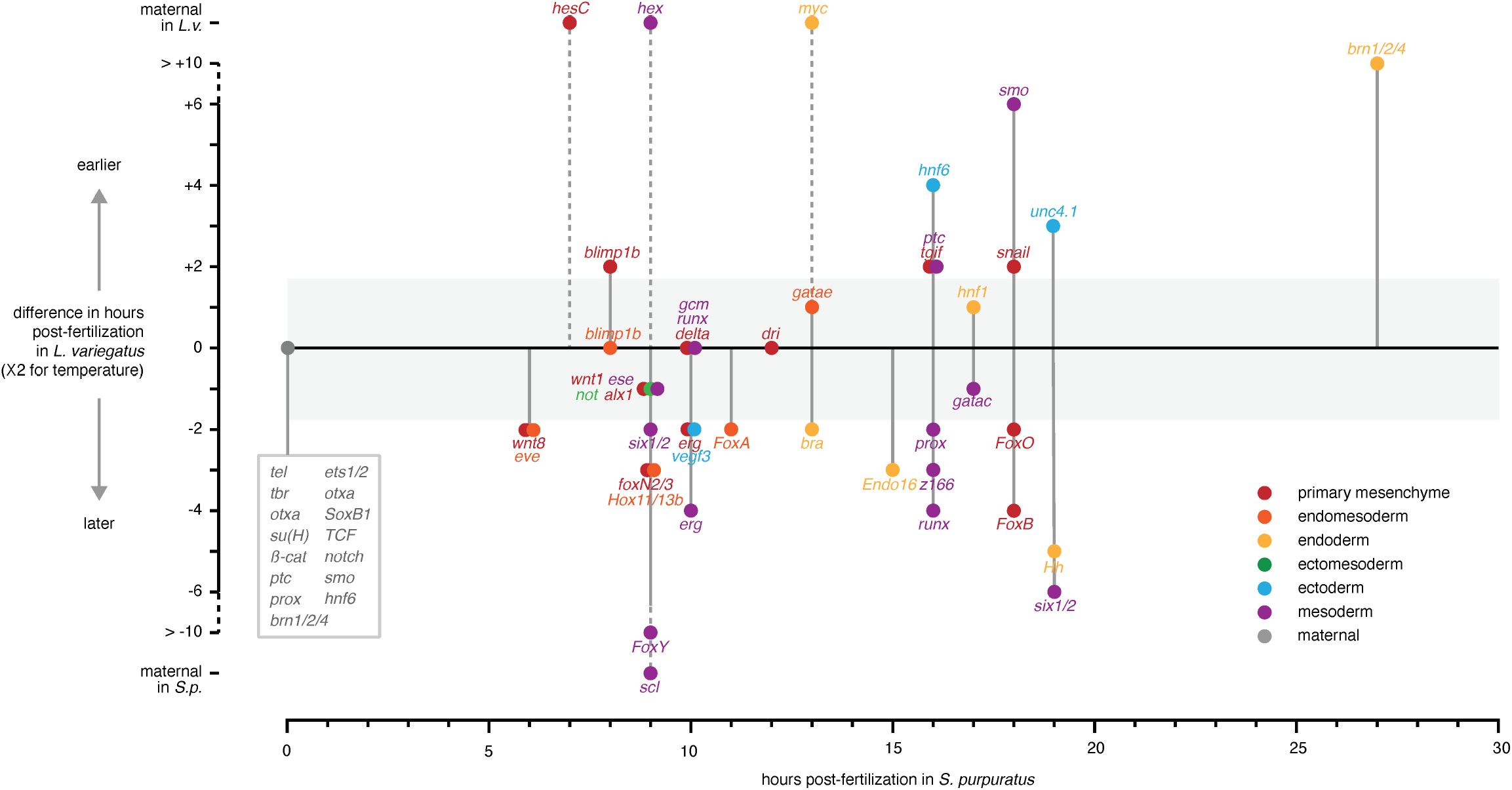
Heterochronic transcriptional activation of dGRN genes. The x-axis indicates the time of first expression in *S. purpuratus,* with each gene indicated by a circle corresponding to germ layer or territory. The y-axis indicates when that same gene is first expressed in *L. variegatus* relative to *S. purpuratus* after normalizing for temperature difference. If a gene is on the horizontal line, its time of first expression is approximately the same in the two species. Due to imprecision in estimating the difference in overall rates of development, differences of <2h are also considered conserved (indicated by the gray box). Genes above the horizontal line and outside the gray box are expressed earlier in *L. variegatus* and those below are expressed later. Many genes are maternally expressed in both species (box pointing to 0 hr), while four are maternally expressed in one species but not in the other *(hesC, hex, myc,* and *scl).* Note that the time of first expression can be uncoupled in different territories (e.g., *blimp1b* is heterochronic in primary mesenchyme but not in endomesoderm). In addition, some genes expressed simultaneously in *S. purpuratus* are expressed both earlier and later in *L. variegatus* (e.g., *smo, snail, FoxA,* and *FoxB* are activated at 19 hpf in *S. purpuratus* but range from 6 hr earlier to 4 hr later in *L. variegatus*). Of the 49 GRN genes plotted here, 15 have conserved timing due to being maternally expressed, 4 have conserved zygotic timing, and 37 have heterochronic zygotic expression in at least one territory of expression.

Of the 49 dGRN genes examined (**Fig. 6** and **Table S5**), 15 are maternally expressed in both species and thus scored as having conserved earliest expression (e.g., *Tel* and *Tbr).* A further five dGRN genes are not maternally expressed and the timing of their earliest zygotic expression is conserved within a particular embryonic territory or cell lineage (e.g., *gcm* and *dri).* However, the majority of dGRN genes (37/49) exhibit heterochronic zygotic first expression within at least one territory of their expression, and in some cases to a substantial degree (e.g., *smo* and *hh).* Of note, the earliest expression of 4 genes shifts between maternal and zygotic *(HesC, hex, myc,* and *scl*). Finally, some dGRN genes show conserved expression within one territory of the embryo but heterochronic expression within another. For instance, the timing of *blimp1* expression is conserved in the endomesoderm but heterochronic in the primary mesenchyme.

Several observations are pertinent. The heterochronies are scattered throughout specification times and differ in sign and magnitude, so there is no “hotspot” or obvious pattern to the temporal shifts. Heterochronies were also observed in each germ layer, indicating that they are spread throughout the dGRN rather than being restricted to a single cell lineage. Note that **Fig. 6** is plotted to show heterochronic shifts of *Lv* relative to *Sp*, but the actual polarity of the evolutionary change is not known (i.e., a given heterochronic shift could have occurred in the lineage leading to either *Lv* or *Sp*). It may be possible to reconstruct the polarity of some shifts based on scRNA-seq datasets from additional sea urchin species. The number of dGRN genes that showed fairly substantial heterochronic shifts in critical regulatory genes raises questions regarding the robustness of dGRNs. With so many evolutionary differences in timing of expression, how is it possible for the dGRNs of these two species to exhibit such an apparently high level of conservation in expression of cell-type specific genes, as well as similar outcomes in response to perturbation experiments? This prompted us to further examine the timing of expression relative to the architecture of the dGRNs in the two species.

### Temporal architecture of the dGRNs

Given the temporal differences we noted in the *Sp* vs *Lv* comparison, we decided to revisit both the *Sp* and *Lv* data and use timing of expression as a driver of the model to see if that had an effect on GRN structure. Our rationale is that timing of expression helps in the analysis of dGRN function. For example, if gene A is perturbed and as a consequence gene B is no longer expressed, the conclusion is that Gene A somehow directly or indirectly drives the expression of Gene B. If timing is considered, however, Gene A could drive Gene B in a forward direction, or, the same perturbation also could be explained by Gene A feeding back to influence expression of Gene B which was expressed earlier, in a direct or indirect feedback mechanism. Experimentally, distinguishing between these two possibilities in constructing circuits often is difficult, but if relative timing of expression of Gene A vs Gene B is known, it is possible to infer directionality of control. Further, since we observed so many heterochronic changes, we wondered whether these would influence feedforward or feedback controls. We thus decided to examine the GRNs of *Sp* and *Lv* to compare the timing of dGRN assembly and explore how timing might influence developmental and evolutionary circuitry.

The temporal modifications to the dGRNs were drawn without changing the results of the earlier perturbation studies in which gene A was shown to regulate gene B. After making the connections, if Gene A, which was known to provide an input into Gene B, was expressed later in time than Gene B, we noted that direct or indirect feedback input with a bold connection line. The most recently published *Sp* dGRN has 12 such feedback inputs. After redrawing all *Sp* and *Lv* dGRNs, (**Fig. S6A-D**) several observations are relevant. The temporally adjusted dGRN models differ significantly in structure from the most recent versions of the *Sp* dGRN (Li et al., 2013; Peter and Davidson, 2015). Of importance, when time of first expression is considered, there is an increase in the number of feedback inputs relative to the original dGRNs. In other words, there are a number of cases (highlighted in the GRN models) in which a gene expressed later has a feedback input into a gene that is expressed earlier. Further, there are substantial differences in the *Sp* and *Lv* dGRNs when the two are compared side by side (**Fig. S6A-D**).

There are about 232 connections in the networks. Not counting inputs in which a gene feeds back on itself, the four dGRNs of *Sp* contain a total of 46 feedback inputs (19.8% of the total number of connections), while the four of *Lv* contain 43 (18.5% of the total). It is well established that feedback can stabilize a network (Brandman and Meyer, 2008), especially negative feedback circuits. As a result of the redrawing of the networks, there are 11 negative feedback inputs in *Sp* and 9 in *Lv* (**Fig. S6)**. Thus it is likely that the capacity for networks to absorb heterochronic changes is at least in part buffered by the presence of these stabilizing feedback loops.

### Gene ontology (GO) enrichment

Much of this analysis has been devoted to examination of the scRNA-seq results in comparison with a large body of knowledge on establishment of dGRNs during development. Many other analyses are possible and promise to be most informative. The database supporting this analysis will be publicly available for anyone with ideas to explore. As one example of where such analyses could lead, analysis of enrichment of gene sets between clusters of cells (**Fig. 1**) can reveal molecular processes that are characteristic of a given cell lineage in the embryo. We used Mann-Whitney U Tests to investigate GO term enrichment within each lineage, based on the percentage of cells expressing a given gene among the most differentially expressed genes within each lineage. The enriched GO terms in some cases match known biological features. For example, ectodermal lineages are highly enriched for expression of genes involved in cilia movement, microtubule processes, and regulation or response to stress. Similarly, clusters composed of cells that are dominated by maternal transcripts are enriched for genes involved in cell cycle processes and regulation of chromatin organization (**Fig. S7**). This analysis also provides insights into additional, less obvious genes that may be involved in distinct cellular functions. For instance, skeletogenic cells are enriched for “regulation of anatomical structure size”, providing potential candidate genes for future experimental analysis aimed at understanding how growth of the larval skeleton is regulated.

## Discussion

The ability to gain new insights through scRNA-seq has been demonstrated in several model embryo systems (Farrell et al., 2018; Foster et al., 2020; Schiebinger et al., 2019; Tintori et al., 2016). Here, we learned that a high quality scRNA-seq analysis detects almost all of the known transcription factors in a well-established dGRN. This is important because dGRNs underlie the specification of all lineages as development progresses, and it is of value to know whether scRNA-seq has the power to assist in further dGRN discovery. Our results demonstrate that once fate specification begins, the sample from each time point contains the expected distribution of cell types, based on published cell lineage analyses. Given that, the 50,000 cells from 18 densely spaced time points enabled us to employ Waddington-OT to trace lineage trajectories and divergences. The optimal transport method was then used to follow temporal profiles of cell fate specification, and to address several unanswered questions related to gene regulation in development. It was of great value to have a well annotated, high quality genome in order to produce gene calls with high fidelity (Davidson et al., 2020). With these quality checks complete, we built on Waddington-OT (Schiebinger et al., 2019) to devise a number of novel computational analyses. Methods were developed to assess timing of first expression of genes within a lineage, and to follow cell fate probabilities based on the optimal transport methods. We had started the analysis with the simple goal of determining how much of the known dGRN was reflected in the scRNA-seq database. When we confirmed that 98% of the genes in the dGRN are detected and readily quantifiable, we then explored the database to assess a number of properties of the dGRN. Further, and looking toward the future, the high degree of concordance between the scRNA-seq database and several comparisons with the dGRNs gave us confidence that the datasets contain, in addition to the known dGRN genes, highly selective candidates for inclusion in future studies of molecular progression toward distinct fates, not only for adding transcription factors, but also for effectors that participate in morphogenesis or a number of biological processes (**Fig. S7**).

Lineage analyses have provided prominent milestones in the advances of developmental biology over the past 120 years. The careful camera lucida tracings of Conklin pioneered descriptions of cell lineages in *Styela* and *Crepidula* (Conklin, 1897, 1905). Innovative transplantation methods were used to trace lymphatic and neural crest origins (LeDouarin and Jotereau, 1973), setting the stage for detailed analyses of how neural crest lineages diverge and contribute to diverse structures in vertebrates. The epic dedication to lineage tracing in *C. elegans* (Sulston and Horvitz, 1977; Sulston et al., 1983), provided the anchor diagram for all lineages and provided the control state for mutational analyses affecting many cell types. More recently, approaches featuring novel methods for fluorescently labeling cells (for example, “brainbow” (Livet, 2007; Pan et al., 2011)), have led to many recent lineage tracing advances.

The optimal transport approach offered by Waddington-OT adds another valuable tool to this endeavor by allowing detailed reconstruction of the sequence of molecular changes that take place within cell lineages. This approach is unbiased, taking into account all genes expressed rather than considering only the transcription factors that characterize the distinct state of a cell at a given time. These transcription factors are represented in the data (**Fig. 4**), and can be tracked to the lineage probabilities, but many other genes contribute to the transcription state of a cell at a given time point during specification and differentiation. Thus, optimal transport offers a powerful new approach for analysis of developmental events that are known to occur at specific times in a lineage, both at the level of gene regulation and at the cellular level in studies of morphogenesis.

The divergence of cells from an uncommitted state to distinct fates is at the core of developmental and stem cell biology. The literature contains a number of examples in model systems where a single molecular event directs a lineage divergence (Driever and Nusslein-Volhard, 1988; Guo and Kemphues, 1995; Kemphues et al., 1988; Nishida and Sawada, 2001; Nusslein-Volhard and Wieschaus, 1980). Those examples, including the skeletogenic cell divergence in the sea urchin embryo, mentioned above (Oliveri et al., 2002; Oliveri et al., 2003; Revilla-i-Domingo et al., 2007), tend to occur early in development. We were curious about the other mode of divergence seen in this study, an asynchronous and an apparent delayed commitment displayed by several of the lineages. This divergence from a common ancestral state appears messy, with many cells seemingly uncommitted to their final fate until quite late (**Figs. 3, 4**). We selected embryos from cultures that were developing as synchronously as possible, so we knew the cells in a given sample were from embryos at the same stage. We examined the specification state of individual cells by determining which dGRN signature genes were expressed in seven intermediate cells randomly chosen from the intermediate zone (**Fig. 5**). These intermediate cells express dGRN signature genes of both fates until late in specification space (the latest we sampled was at 16 hpf which corresponds to the end of gastrulation in *Lv*). The transcriptional state of these cells moves quite far from the ancestral state over time, as seen by the progress across the triangles that represent specification space (**Figs. 3-5**), and yet they remain uncommitted, at least based upon the probabilities calculated from all genes expressed. This delayed specification of some cells could be quite valuable for an embryo, for instance by contributing to flexibility in the exact number of fully differentiated cells of distinct fates. Whether this actually the case will require further study.

The published sea urchin dGRN was experimentally established in detail and repeated by many laboratories over two decades, providing a strong set of priors to assess. The data here do not challenge, nor are they capable of challenging, results of the perturbation experiments that were used to establish the connections in the published GRNs. Rather, our analyses assess the ability of scRNA-seq to reflect the spatiotemporal expression of the genes within the dGRN. We asked how predictive might the scRNA-seq approach be in producing a provisional dGRN? At any given time point, a pick of the top 200 genes in a lineage (based on a ratio comparison with expression in cells of the “other” lineage), included about 72% of the genes depicted in dGRN models of that tissue at that time. The remaining genes were also present in the scRNA-seq lineage time points but not in the top 200 lists because they either were expressed ubiquitously, or also expressed strongly in other lineages, thereby reducing the ratio. We conclude that the optimal transport approach operationalized by Waddington-OT offers an efficient way to identify candidate transcription factors in a given cell lineage prior to carrying out perturbation studies.

Experiments leading to the assembly of the *Sp* and *Lv* dGRN models have revealed a remarkable degree of conservation despite the 40-50 million years that have elapsed since their last common ancestor (McClay et al., 2020). At an even greater temporal distance of 500 million years, sea urchins and sea stars also share some conserved network circuits, although some transcription factors have assumed different roles in cell fate specification (Hinman and Davidson, 2007). Thus, over vast periods of time, networks are amenable to evolutionary rewiring, but the architecture appears constrained and resists change, as reflected by the small number of differences in dGRN models between *Sp* and *Lv*. Those models, however, did not include a careful analysis of timing of gene expression. They are also somewhat limited in that they consider transcription factors known at the time, and are not aware of other transcription factors and signals that may have important roles that remain uncharacterized. A dense scRNA-seq time course can amplify candidate gene identification, leading to deeper insight into the mechanisms of face specification in embryos. Our finding that specification within some cell lineages occurs asynchronously and often quite delayed suggests we have much to learn of how the cells of an embryo, especially a regulative embryo like the sea urchin, move toward their respective fates.

## Methods

### Embryo Spawning and Culture

Six female urchins were spawned by injecting 1ml 0.5M KCl intracoelomically. Unfertilized eggs were allowed to settle and washed three times in artificial sea water (ASW). Eggs were then resuspended, and fertilized by a single male’s sperm in .03g PABA/100 ml ASW. Following fertilization, eggs were washed three additional times in ASW to remove residual PABA. The fertilized embryos were then combined and co-cultured together at 22-23 degrees Celsius while gently being stirred by a motorized stir rod. Embryos were then sampled at various time points to be dissociated and fixed for scRNA-seq. At each stage the embryos collected were very similar in stage to each other, an important consideration for temporally following development of the cells in this study.

### Embryo Dissociation and Fixation

Once embryos developed to the appropriate stage, a portion of the co-culture was taken, and washed two times in Calcium-Free Artificial Seawater (CFASW). After washing embryos with CFASW, they were dissociated by gentle trituration after 10 minutes in dissociation buffer, made of 1.0M Glycine and 0.25mM EDTA, pH 8.0 with HCl at 4 degrees Celsius, and on the rocker. Following dissociation, cells were resuspended in CFASW, and fixed at a final concentration of 80% Methanol in CFASW for one hour, at 4 degrees Celsius. Following fixation, cells were stored at −20 degrees Celsius, and all cell libraries were processed within one month of dissociation.

### Rehydration of Methanol Fixed Single Cells for Library Preparation and Sequencing

Following fixation, cells were washed twice, and rehydrated in a 3x Saline Sodium Citrate buffer before cell count and library preparation. 19 single cell libraries were prepared using the 10x Genomics 3’ v3 gene expression kit and the 10x Chromium platform to encapsulate single cells within droplets. Library quality was verified using the Agilent 2100 Bioanalyzer. Libraries were pooled and Duke Genomics and Computational Biology Core facility sequenced samples across two NovaSeq6000 S1 flow cells with 28 x 8 x 91 bp sequencing performed.

### Computational Analysis

#### Data download, FastQ file generation, Genome Indexing, Genome Mapping and Counting, production of raw csv counts files

Following sequencing, we used Cellranger 3.1.0 to convert Illumina-generated BCL files to fastq files using the Cellranger “mkfastq” command. We then applied the “mkref’ command to index the most recent *Lv*3.0 Genome (Davidson et al., 2020). We then used the “count” command to demultiplex and count reads mapping to the reference *Lv* genome. The “mat2csv” command was used to get CSV RNA count matrix files for each sample for further downstream analysis. In addition, we used the command “aggr” on all samples to generate an automated 10x Cloupe browser that is easily accessible, and requires no coding experience to utilize.

#### Filtering and normalization

All CSV RNA count matrix files were uploaded to R and a seurat object was generated for each sample. All seurat objects were then merged to undergo uniform quality control, normalization and data exploration with all 19 samples. The merged object was then filtered to remove low quality cells with nFeature_RNA > 200, nFeature_RNA < 7500, and percent.mt < 5. SCTransform was then applied to the merged filtered object to perform normalization and removal of technical variation, while preserving biological variation and various processes (see below). The data was then scaled, and variable features amongst the cells were found.

#### Dimensionality reduction, visualization, and clustering

We next performed Principal Component Analysis on the SCTransformed seurat object file, and found the nearest neighbors using 105 PC dimensions of variable gene space. UMAP was applied to multi-dimensional scRNA-seq data to visualize the cells in a two-dimensional space. Finally, clustering was performed using graph-based Louvain Clustering with resolution, res=2.4, resulting in 63 clusters. The 63 clusters were annotated using dGRN genes, and published in situ hybridization patterns as markers.

#### Inferring developmental trajectories with Waddington-OT

We next applied Waddington-OT to infer developmental trajectories. As input, we used the SCTransform normalized expression matrix together with expansion rates estimated from expected changes in proportion of lineages over time (**Fig. S2**). Growth rates were estimated by lineage based on the expected number of cells of each lineage at key developmental time points. Growth rates were assumed to be uniform between estimates of expected number of cells. Maternal growth rates were assumed to be the expected growth rate across all cells at that time point.

Two additional growth rates were fit based on two cell cycle scores using a sigmoid function to smoothly fit growth rates between the minimum and maximum expected at that time point. Validation values were very similar between the two growth rates fit on the cell cycle scores and the growth rate based on expectation alone. For simplicity, the expectation-based growth rate was used throughout the analysis.

Transport maps were calculated using the growth rates described above, the optimization parameters *ε* **=** 0.05, **λ**1= 1, and **λ**2 = 50, and a single iteration of growth rate learning.

#### Validating trajectories with geodesic interpolation

We validated our results by demonstrating that we can interpolate the distribution of cells at held-out timepoints (**Fig. S4**). For each triplet of consecutive time-points (e.g. 5,6,7 or 6,7,8 etc.), we held out the data from the middle time-point and attempted to reconstruct it by connecting the first to the third. We then quantified our performance by comparing to the held-out midpoint.

The blue curve shows the results from optimal transport, which is lower than various null models (yellow, orange, green, purple). The null models are:

- “Random” (Orange): we randomly connect cells to descendants.
- “Random with growth” (yellow): We randomly connect cells to descendants, incorporating the same estimate of growth as for OT.
- “First” (Green): We use the first time-point in the triplet to estimate the second element of the triplet.
- “Last” (Purple): We use the third element of the triplet to estimate the second element.

In order to test the stability of our results, we varied parameters of optimal transport over an order of magnitude in each direction (see **Fig. S4B-D**). Additionally, we repeated the analysis on datasets with downsampled cells and reads as low as 10% of cells and 500 UMI per cell respectively. Downsampling cells, we found, using Waddington-OT, outperformed null methods for all proportions and only saw a gradual increase in validation values (**Fig. S4E**). Downsampling reads, we found Waddington-OT outperformed null methods down to 500 UMI (**Fig. S4F**).

#### Visualizing divergence of fates with barycentric coordinates

In order to visualize the divergence of fates, we developed a simple way to visualize fate probabilities for triples (or quadruples) of lineages in a triangular (or tetrahedral) plot. The visualization employs barycentric coordinates to represent k-dimensional probability vectors in k-1-dimensional space. We identify a corner of the triangle for each of these possible fates, and position the cells according to their relative probabilities as follows:

Let *a, b, c* denote the vertices of the triangle in *R^2^* and let *p* = (*p*_1_, *p*_2_, *p*_3_) denote the probability vector we wish to visualize. The components of *p* are used as coefficients in a convex combination of the vertices. In other words, the probability vector *p* is mapped to *p*_1_,*a* + *p*_2_*b* + *p*_3_*c* ∈ *R^2^*. Note that *p*_1_, + *p*_2_ + *p*_3_ = 1, so each probability vector is mapped to a point inside the triangle.

Cells perfectly fated to obtain a single fate are positioned exactly at the corresponding vertex, while cells with indeterminate fates are positioned in the interior of the triangle. The very center of the triangle corresponds to cells that are equally likely to choose any of the three fates, and cells along an edge have zero chance of reaching the opposite vertex. **Fig. 3** illustrates examples of how selected pairs of lineages diverge in *Lv* over time. **Movie S1** shows a tetrahedron visualization of the divergence of four fates simultaneously.

#### Visualizing Gene Regulatory Networks

We visualized signature genes from the dGRNs on the triangle plots and the UMAP plots using GRN score ratios. The illustration aimed to show how intermediate cells tend to express networks from both possible fates. GRN score was defined as the fraction of genes from the signature list that are expressed in a cell. Each cell was given two of these scores, each corresponding to a GRN of interest. To compare the expression of one regulatory network to the other, we defined a GRN score ratio as 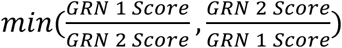. The ratio produces values between 0 and 1. A value of 0 means only one of the networks is expressed in the cell. Meanwhile, a value of 1 means both networks are expressed in equal proportion. **Figs. 5 and S5** shows cells colored by this ratio.

### Gene Ontology Analysis

We used functional enrichment analyses to determine if there are any biological processes (BP) and molecular functions (MF) that were overrepresented among each lineage based on differentially expressed genes. Enrichment analyses were conducted with a rank-based gene enrichment method that was originally implemented for bulk-RNAseq data, GO-MWU (Wright et al., 2015). Here, we adapted this method to process data output from our single-cell data output files of differential gene expression where we used pct.1 (percentage of cells with differentially expressed gene) instead of kME values (module membership scores). In this case, we implemented global Fisher’s exact test for presence-absence of functional categories in the cluster, and then, within cluster MWU test for association of the included categories with higher pct.1 values. Results were plotted using alluvial plots, here we plotted the top 5 BP and MF categories in each lineage according to their lowest p-adjusted values and used custom R scripts to consolidate data tables and plot figures (**Fig. S6**).

## Acknowledgements

The authors thank members of the McClay, Wray and Schiebinger labs for their critical input and feedback. We also appreciate the help provided by Nicolas Devos, Ph.D, and the Duke GCB core facility. We also appreciate the assistance provided by Philip Benfey, Ph.D lab members, especially postdoctoral fellows Rachel Shahan, Ph.D and Isaiah Taylor, Ph.D in the Biology Department.

